# SwarmSight: Real-Time Tracking of Insect Antenna Movements and Proboscis Extension Reflex using a Common Preparation and Conventional Hardware

**DOI:** 10.1101/183459

**Authors:** Justas Birgiolas, Christopher M. Jernigan, Richard C. Gerkin, Brian H. Smith, Sharon M. Crook

## Abstract

Many scientifically and agriculturally important insects use antennae to detect the presence of volatile chemical compounds and extend their proboscis during feeding. The ability to rapidly obtain high-resolution measurements of natural antenna and proboscis movements and assess how they change in response to chemical, developmental, and genetic manipulations can aid the understanding of insect behavior. By extending our previous work on assessing aggregate insect swarm or animal group movements from natural and laboratory videos using video analysis software SwarmSight, we developed a novel, free, and open-source software module, SwarmSight Appendage Tracking (SwarmSight.org) for frame-by-frame tracking of insect antenna and proboscis positions from conventional web camera videos using conventional computers. The software processes frames about 120 times faster than humans, performs at better than human accuracy, and, using 30 frames-per-second videos, can capture antennal dynamics up to 15 Hz. We used the software to track the antennal response of honey bees to two odors and found significant mean antennal retractions away from the odor source about 1 s after odor presentation. We observed antenna position density heat map cluster formation and cluster and mean angle dependence on odor concentration.

## INTRODUCTION

Most arthropods move antennae or other appendage to sample environmental cues and signals in time and space. The animals can use the antennae to navigate their environment by detecting sensory cues such as chemical volatiles and gustatory and mechanical stimuli^1–4^. In insects, the antennae contain sensory receptors that bind to chemical volatiles^4–6^ and transmit this signal via olfactory sensory neurons to central brain regions^1,7–9^. The insects can adjust antennae positions to modulate information about incoming odors^4,10,11^. This modulation facilitates actively informed behavioral responses to odors and their plumes^12,13^.

Many insects, including Hymenopterans (e.g. honey bees and bumblebees), Lepidopterans (e.g. butterflies), and Dipterans (e.g. flies and mosquitoes), among others, feed by extending their proboscis^14–21^. Proboscis extension has been reliably used in the past for a variety of learning and memory tasks^22–31^. Similarly, quantitative assessment of antennae movement with high temporal and spatial resolution might yield insight into the relationship between the stimulus, the behavior, and internal state of the animal. Indeed previous work has shown how the antennal movements contain a rich amount of information about honey bee tracking of the environment and how the movements change with learning^32–38^.

In the last decade, methods for observing animal behavior have been greatly accelerated by advances in high-resolution video cameras, computer processing speeds, and machine vision algorithms. Tasks like animal detection, counting, tracking, and place preference analyses have been aided with sophisticated software that can process videos of animal behavior and extract relevant measures^39–47^.

These technologies have also aided tracking of insect antenna and proboscis movements. It is possible for human raters to use a mouse cursor to manually track the position of the antennae. However, while this method can be accurate, the task is time consuming, and human inattention and fatigue can result in unreliable results. Special equipment and preparation can be used to reduce the need for complex software. For example, one setup used a high-speed camera and painted the tips of the antennae to track the antenna movement^48^. Users can also be asked to select key-frames of videos to assist the software in detecting the antenna and proboscis location^49^. Another approach detected the two largest motion clusters to identify antennae, but it does not detect the proboscis location^50^. Another software package can detect antenna and proboscis locations, but requires about 7.5 seconds of processing time per frame^51^, which could be prohibitive for real-time or long-term observation studies. Finally, it might be possible to customize commercial software packages like EthoVision to perform the task^46^, but their licensing and training costs can be prohibitive.

With the method described here, we extended our previous work on motion analysis software^41^ to track the locations of insect antennae and proboscis with the following goals: (1) no requirement for special hardware or complex animal preparation, (2) frame processing at real-time (30 frames per second or faster) on a conventional computer, (3) ease of use, and (4) open-source, easily extendable code.

The resulting novel method and open-source software, *SwarmSight Appendage Tracking*, does not require painting of the antennae tips, can use a consumer web camera to capture videos, and processes video frames 30-60 frames per second on a conventional computer (Figure 1). The software takes video files as input. The user locates the position of the insect head in the video and, after processing, a comma separated values (.csv) file is produced with the locations of the antennae and proboscis. The software is capable of reading hundreds of different video formats (including formats produced by most digital cameras) through the use of the FFmpeg library^52^.

**Figure 1:**
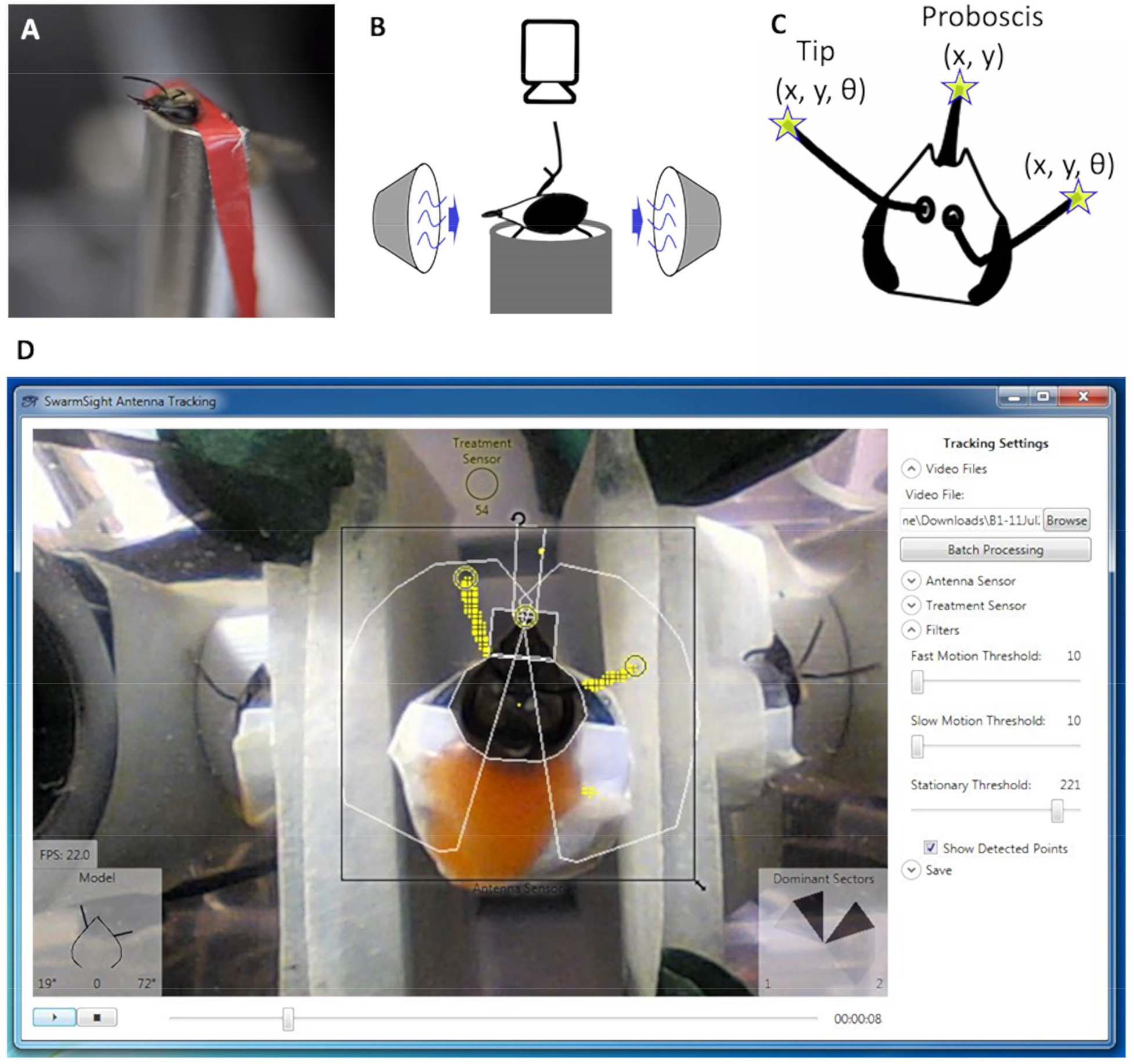
Animal Setup and Software Output. A) A honey bee forager with its head and body restrained in a harness. B) Odor source is placed in front of the animal, a video camera is positioned above, and a vacuum source is placed behind the animal. C) The antenna tip and proboscis variables detected by the SwarmSight software from the video. D) The user positions the antenna sensor over the animal and adjusts the filter parameters. The software detects the antenna and proboscis positions (yellow rings).

First, an insect’s body and its head are restrained in a harness such that the antenna and proboscis movements are easily observed (Figure 1A). An odor source is placed in front of the insect, with a vacuum source placed behind, to remove the odors from the air and minimize potential effects of sensory adaptation (Figure 1B). A conventional web camera is placed above the insect’s head on a tripod. An LED can be positioned within the camera’s view to indicate when the odor is being presented.

After filming, the video file is opened with the SwarmSight software, where the user positions the Antenna Sensor widget (Figure 1D, black square) over the head of the insect, and starts the video playback. When the results are saved, the .csv file will contain the X, Y positions of the antenna tips, the antenna angles relative to the front of the head (Figure 2), and the proboscis X, Y position. Additionally, a dominant sector metric is computed for each antenna. The metric shows which of the five 36-degree sectors surrounding each antenna contained the most points deemed likely to be the antennae, and can be useful if the antenna position/angle metrics are not reliable due to noisy or otherwise problematic videos.

**Figure 2:**
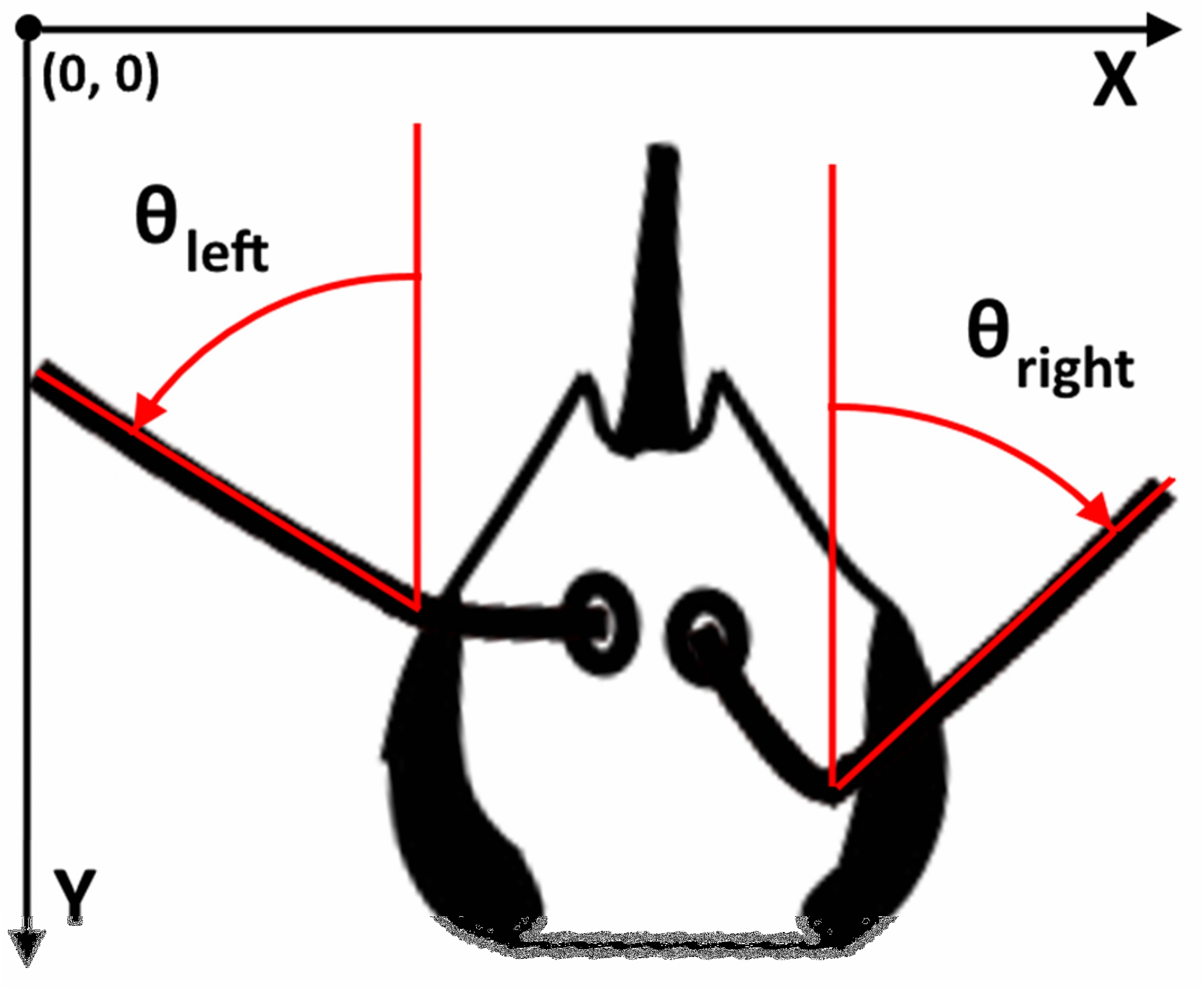
Antenna Coordinate System. X, Y values use the video coordinate system, where top left corner is the origin and X and Y values increase when moving towards the bottom right corner. Angles are expressed in degrees with respect to the front of the head (usually the odor source). A “0” value signifies that the line formed by the antenna flagellum is pointing directly in front of the animal. All angles are positive, except when an antenna points to the opposing direction (e.g. right flagellum points to the left).

Briefly, the software works by using a set of motion filters^53^ and a relaxed flood fill algorithm^54^. To find likely antenna points, two filters are used: a 3-consecutive-frame difference filter^41,55^ and a median-background subtraction^56^ filter. A color distance threshold filter is used for proboscis point detection. The top 10% of the points of each filter are combined, and a flood fill algorithm that inspects contiguous points with gaps up to 2 pixels locates extreme points. Parallel frame decoding, processing, and rendering pipelines and optimized memory allocation of the filter data flow achieves high performance. The raw x and y coordinate values produced by the software are post-processed with a 3-frame rolling median filter^57^ (see Discussion). The instructions to download the full source code can be found online^58^.

Below, we describe a protocol to prepare a honey bee forager for antenna tracking. A similar protocol could be used to track the antenna/proboscis movements of any other insect. In the results section, we describe a sample antenna trace output that is detected by the software, the comparison of the software output to tracking performed by human raters, and assessment of antennae movement in response to five odorants.

### PROTOCOL

1. Catch and harness honey bees: follow the Protocol steps 1 through 3.1.1 of Smith & Burden (2014)^59^.

2. Preparing the Animal Harness and Video Camera

2.1. Hide the legs by applying tape over the top of the harness tube, visually inspecting that legs cannot be seen moving from the top.

2.2. Restrain the head by applying heated wax to the back of the insect head. Visually inspect that the head is fixed and not moving. At this point, the antennae and the mandibles should be the only appendages free to move.

2.3. Maximize contrast between the antennae and the video background by placing a white sheet of paper underneath the insect harness. Minimize the need to later adjust the camera by marking the location of the insect harness on the paper, and then placing new individuals at the same location.

2.4. Fix the camera position using a tripod to place the camera above the insect’s head. Using the camera software, preview the video, and zoom in to magnify the head image, allowing for a ~20-30% clearance on all sides of the video.

2.4.1. Ensure that the only moving objects in the camera view are the antennae or the proboscis/mandibles and reposition the camera or the animal if necessary.

Note: SwarmSight checks for movement in pixels surrounding the head. Extraneous motion in the immediate vicinity of the head caused by objects such as legs, shadows, fans, or humans may confuse the software, and introduce additional noise.

2.5. Minimize antenna shadows by adjusting ambient lighting. The software can tolerate some shadows, but for best results, they should be kept to a minimum.

2.6. Prevent automatic camera exposure adjustments by using camera shutter speed software to keep camera exposure time constant throughout the video. Using the software, adjust the shutter speed to maximize contrast (video scene not too light or too dark).

Note: In ArcSoft WebCam Companion (comes with the Genius WideCam F100 camera, see Materials), adjust the Exposure slider under Webcam Settings, Advanced. In iGlasses, adjust Exposure Time slider under Basic tab.

2.7. Place odor delivery source and ensure that it does not obstruct the camera view by inspecting the camera video feed. Ensure that a vacuum source is placed on the opposite side to remove the stimulus odors.

2.8. Place an LED, or some other visual indicator that changes brightness to indicate odor delivery, within the camera view. The LED brightness value is saved by the software and can be used to determine the exact frames when the odor delivery begins and ends.

3. Film each Individual under Experimental Conditions

3.1. Film each individual insect and test condition in separate video files by either recording each individual-test combination separately or using video editing software to split a long video file into smaller files.

Note: The software requires the user to locate the position of the head in each video, and for the head to remain fixed. If the head moves, additional noise will be introduced. The Batch Processing feature of SwarmSight allows the user to rapidly set the location of the head for multiple videos and assumes that the insect head remains fixed for the duration of each video file. Instructions how to split long video files can be found online^60^.

4. Video Analysis

4.1. Download and install the Antenna Tracking module of SwarmSight by following the steps provided online^58^.

Note: Video tutorials describing how to use the software are available on the website as well.

4.2. Open a video file showing a filmed animal by using the Browse button.

4.3. Positioning the Antenna Sensor and Treatment Sensor

4.3.1. Once the video loads, position the rectangular “Antenna Sensor” widget over the animal’s head, using the rotation and scale icons to align the widget with the head (see Figure 1D for example).

4.3.2. Position the circular “Treatment Sensor” widget over the LED that indicates when the odor or stimulus is being presented. The Treatment Sensor will record the brightness value of the pixel at the center of the widget for every frame.

4.4. Starting Video Processing

4.4.1. Press the “Play” button (black triangle) in the bottom left corner to start the analysis of the frames.

Note: The detected likely antenna and proboscis points will be highlighted yellow. The yellow rings will show the location of the tips of the appendages. The angles (where 0 is directly in front of the animal) of the antenna and the proboscis extension length will be shown in the “Model” widget in the lower left corner (see Figure 1D). The “Dominant Sectors” widget in the lower right corner will show the relative intensity of the five 36-degree sectors where the most antenna points have been detected. The darkest sectors contain the most points, while the lightest have the fewest. The sector number (1-5) with the most points will be shown in the lower corners of the widget (see Figure 1D).

4.5. Adjusting Filter Thresholds and Adding Exclusion Zones

4.5.1. To change the sensitivity of the filters, adjust sliders in the “Filters” section, on the right panel.

Note: Depending on the lighting conditions and general movement speed of the appendages, different filter sensitivities will be optimal. The user can find the optimal values by adjusting the values and observing the highlighted areas in the Antenna Sensor widget. When an ideal set of sensitivities is found, only the appendages will be highlighted. It is recommended to fast forward to other parts of the video to ensure the filter sensitivities are optimal there, too.

4.5.2. Optionally, to ignore extraneous objects, on the right panel, expand the “Antenna Sensor” section, click the “Add Exclusion Zone” button (see Figure 1D), and click on a set of points to form a red polygon, the contents of which will be ignored by the software.

Note: If the video contains extraneous motion, and the motion is within the Antenna Sensor widget’s zone (e.g. moving legs, strong shadows, lab equipment, etc…), the software may mistake it for appendage movement. The extraneous objects can be ignored by drawing red polygons or “Exclusion Zones”. Anything inside a red polygon will not be used for tracking.

4.6. Saving Results

4.6.1. Once the filters and widgets have been set up, stop the video, restart it from the beginning, and play it to the end.

Note: Once the whole video has played, the positions of the appendages for all video frames will be stored in memory.

4.6.2. To save the appendage position data to a file, expand the “Save” section on the right, and click the “Save to .CSV” button. Then chose a folder where to save the file.

Note: The “Save to .CSV” button will save the processing results to a .csv file. By default, the user will be offered to save the .csv file in the same folder as the video file and will have a date and time as part of the file name. The resulting .csv file will contain a set of columns that contain information about the position of the appendages, including the antenna angles and dominant sectors, as well as orientation and position of the head. The description of each column is provided online^61^.

4.6.3. Optionally, use the Columns(s) and Value(s) fields in the Save section to create an extra column (or more if separated by commas) in the .csv file to record information such as subject ID or the name of an experimental condition. The value in the Column(s) box will appear in the header of the first column and the value in the Value(s) box will be repeated in all rows of the first column.

4.7. Batch Processing

Note: The software can process multiple video files in a batch. However, the user must provide the head location information for each video before starting the batch.

4.7.1. In the right panel, in “Video Files” section, click the “Batch Processing” button to open a window that allows creating a list of video files to be processed sequentially by the software.

4.7.2. Use the “Add More Video Files to Batch” button to select one or more video files to be included in the batch list.

4.7.3. Optionally, use the “CTRL” or “SHIFT” keys to select multiple videos that will use the same set of widget parameters.

Note: Good candidates for parameter reuse are sets of videos of the same animal that has not been moved between different experimental conditions.

4.7.4. Start setting the widget parameters to be used for the selected videos by clicking the “Set Sensor Positions for Selected” button.

4.7.5. Adjust parameters in the Antenna Sensor, Treatment Sensor, Filters, or Save sections, and click “Save Parameters to Batch” when done.

4.7.6. Once the parameters for each video have been selected, start the batch process by clicking the “Start Processing” button.

Note: The software will load the video files, in the order in which they appear in the batch list, process them, and save their corresponding .csv files to the same folder where the video files are located. A progress bar at the top will provide an estimated finish time after the first video has been completed.

### REPRESENTATIVE RESULTS

In the sections below, we describe an example plot of antennae angles produced from the data of the software, comparison of the software accuracy and speed with human raters, and the results of an experiment where honey bee antenna movement is affected by presentation of different odors. R software^62,63^ was used to perform the analysis and generate the figures. R code for analysis and figure generation as well as video tutorials can be found online^58^.

### Software Output

Figure 3 shows five randomly selected traces of antenna angles detected by the software from videos of honey bees presented with pure and 35x mineral oil diluted versions of heptanal and heptanol, as well as clean air.

**Figure 3:**
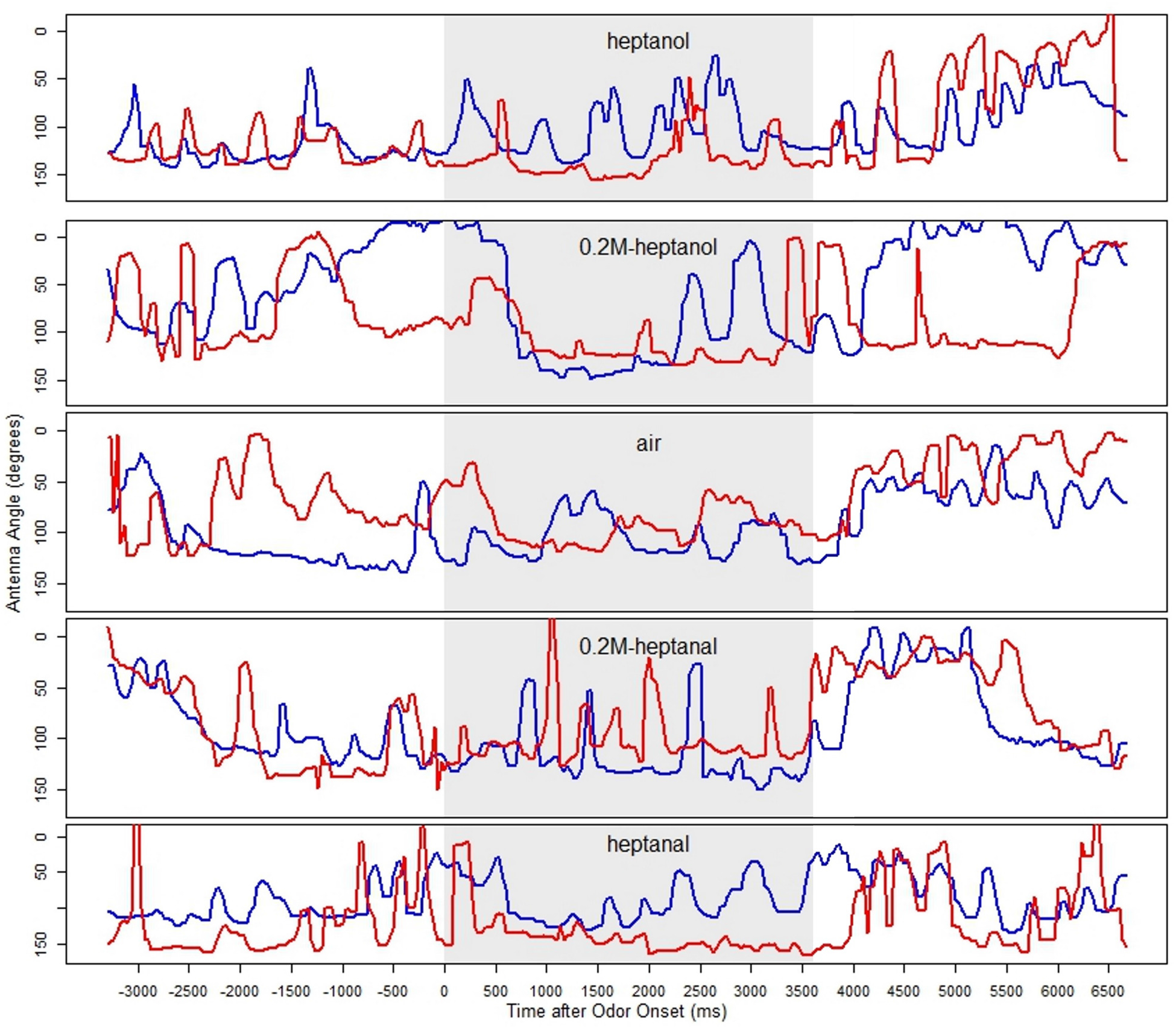
Five Sample Traces of Antennae Angles Detected by SwarmSight. Y-axis shows antenna angle in degrees, where “0” is directly in front of the animal, towards the odor source, with larger values pointing away from the odor source. Heptanol, heptanal, and their 35x mineral oil diluted versions, as well as clean air, were applied during the gray 0-3600 ms windows to single honey bee foragers. Left antenna is marked red, right marked blue. Five random bees, one from each condition, are depicted in the five plots.

### Software Validation

To validate that the software can reliably detect the locations of the antennae, we compared human-located antenna positions with the positions located by the software. Two human raters were asked to locate the antenna and proboscis tips in 425 video frames (~ 14 s of video). A custom software module recorded the appendage locations marked by the raters, automatically advanced video frames, and recorded the amount of time spent on the task. As an example of correspondence between human- and software-located values, superimposed vertical coordinate traces of one antenna for the software and for the two human detected locations are shown in Figure 4A. The distance between the two raters’ marked antenna positions was computed and named “Inter-Human Distance”. The distance between the antenna location detected by the software and the closest location detected by the human raters was computed and named “Software-Closest Human Distance” (Figure 4B).

**Figure 4:**
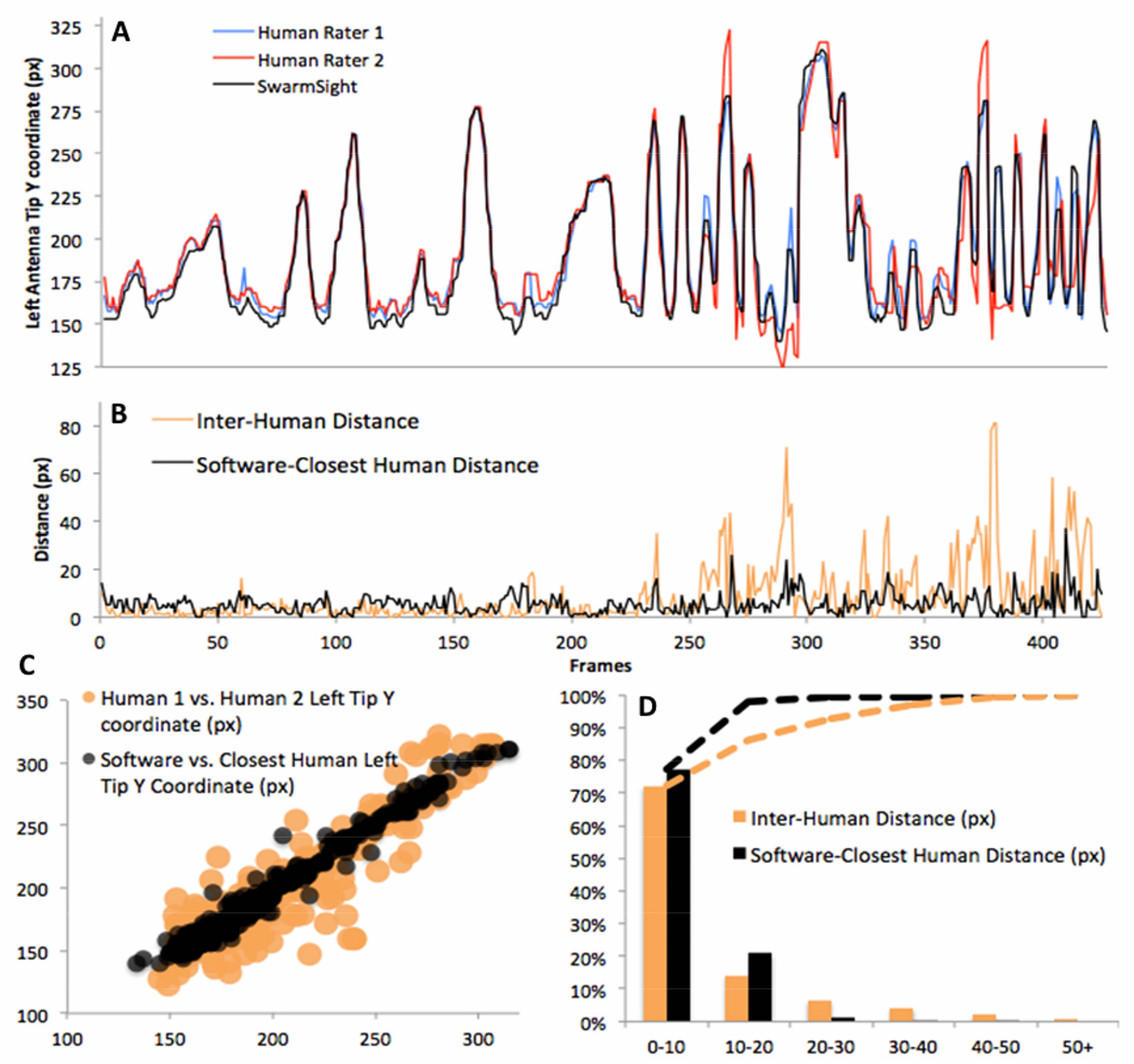
Comparison with Human Raters. A) Two human raters and SwarmSight located antenna tips in 425 video frames. The frame-by-frame left antenna tip Y coordinates found by the human raters and software are superimposed. B) Superimposed frame-by-frame disagreement (distance in video pixels) between human raters (orange) and disagreement between software and closest human rater value (black). C) Human vs. human antenna tip locations (orange) and software vs. human locations (black). D) Histograms and cumulative distributions (dashed) of human vs. human and software vs. human frame-by-frame disagreement distances.

Inter-Human Distance was 10.9 pixels (px) on average, within 55.2 px in 95% of the frames, and had a maximum value of 81.6 px. The Software-Closest Human Distance was 8.0 px on average, within 18.3 px in 95% of the frames, and had a maximum value of 49.0 px (see distance histograms in Figure 4D, and Figure 4C). 5 px was approximately the width of an antenna. Overall, the Inter-Human Distance was small for the frames at the beginning of the task, and increased in the second half of the task. We suspect this was due to rater fatigue. Meanwhile, Software-Closest Human Distance levels remained constant throughout the task.

### Software Speed and Accuracy Comparison with Human Raters

Humans rated antenna tip and proboscis locations at an average speed of 0.52 frames per second (fps). To estimate human fps, the total number of frames rated by humans (425 each) was divided by the total time they spent on the task (873s and 761s). The software rated the frames at 65 fps on average on a Dual-Core Windows 7 PC. Together with high processing speed and accuracy similar to or better than human raters, the software can be expected to perform the work of about 125 human raters per unit of time.

### Detection of Antenna Response to Odors

To demonstrate that the protocol can be used to detect significant behavioral differences in insect movement, we subjected 23 female honey bees to two different odors. Pure heptanal and heptanol, 35x mineral oil dilutions of the two odors, and clean air as the control, were presented each for 4 s (five conditions in total). Videos, as described in the protocol above, were processed with SwarmSight software, and the antenna angles analyzed (Figure 5).

**Figure 5:**
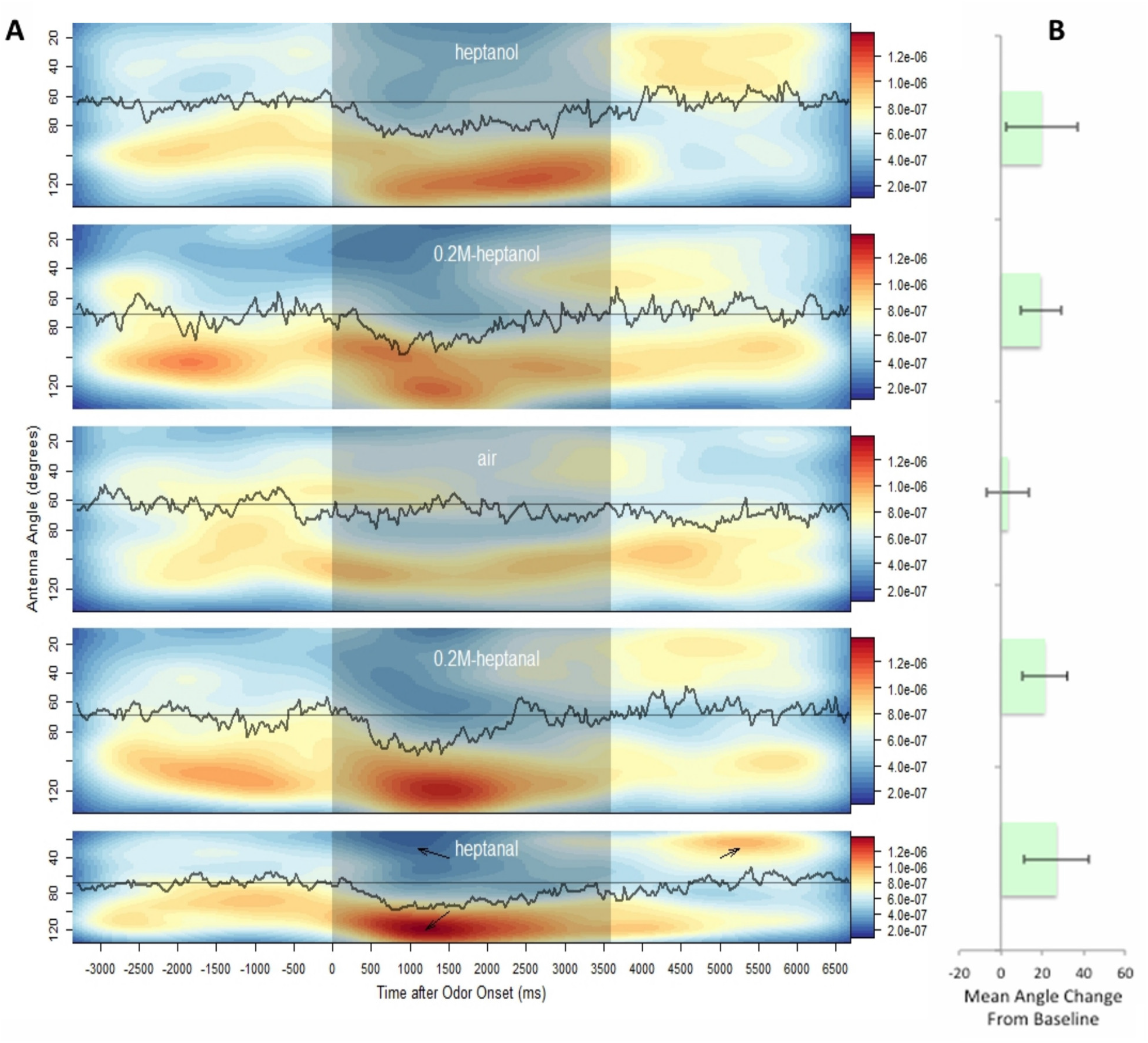
Antenna Angle Means and Density Heat Maps for Five Odor Conditions. A) Heat maps showing antenna angle density before, during (darker middle region), and after administration of heptanol, air, and heptanal odorants to female honey bees (n = 23). Black curves are per-frame average antenna angles (both antennae). Horizontal lines are pre-odor mean (baseline) angles. Note the cluster of preferred antenna locations (red cluster in bottom plot) away from the odor source for pure odor conditions, and corresponding changes to the mean antenna angle. Also note the “rebound” cluster after odor conclusion and its apparent onset dependence on odor concentration (see cluster location in the other four plots). Density heat map color scale is arbitrary but uniform across all conditions. B) Mean angle change from pre-odor mean (error bars S.E.M). Except for air, all mean changes were significant (t-test p < 0.05).

Video frames from 9 s segments of the video consisting of 3 s before the odor onset, 3.6 s of odor presentation, and 2.4 s after odor conclusion were aligned across all individuals and conditions (300 frames per segment). The per-frame means of both antenna angles of all individuals were computed for each condition and called “Mean Angles” (Figure 5A, black curves). The mean antenna angles of frames before the odor onset across individuals for each condition were computed and called “Pre-Odor Baselines” (Figure 5A, thin horizontal lines).

In all conditions, except control, the mean angles increased from baselines, each peaking once 750-1050ms after the odor onset (Figure 5A, black curves in 0-3600 ms region). The mean changes from baselines were tested for significance (Figure 5B) by comparing the two-antenna means of individuals at the peak odor-presentation mean angle time of each condition to the baseline mean using a series of 1-sample t-tests (Shapiro normality tests N.S. in all conditions). The mean angle change from baseline was 26.9° for pure heptanal (mean peaked at 750 ms after odor onset), 21.1° for 0.2M heptanal (at 990 ms), 19.6° for pure heptanol (at 1050 ms), 19.3° for 0.2M heptanol (at 780 ms), and 3.45° for air control (no peak). In all conditions, except control, the mean angle change from baseline was significant (Holm adjusted p < 0.05). We note that the mean angle takes longer to return to baseline in response to pure odorants than to diluted odorants (low-pass filtered mean returned to baseline 3,690 ms after odor onset for pure and at 2,940ms for diluted heptanol; for heptanal, return times were 4,260 ms for pure and 3,000 ms for diluted versions).

### Visualization using Heat Maps

To visualize the antenna responses, antenna angle density heat maps for each condition were generated (Figure 5A, blue-red background). Antenna angles across the 10 s video segments for each individual per condition were convolved with a Gaussian kernel (R package MASS, kde2d function^64^). Blue areas show low densities of antenna angles, while red areas show high densities of antenna angles. The heat map in the bottom plot of Figure 5A for the pure heptanal condition illustrates the antenna behavior.

The map shows that before the odor is presented (t < 0), the antenna angle density is distributed relatively uniformly across all angles. About 1 s after odor onset (t ~ 1000 ms), a pair of blue and red clusters appears. In areas shaded red, the antennae were found more frequently than in areas shaded blue. The blue cluster indicates that antennae tended to avoid smaller angles (odor source was located in the direction of 0 degrees), while the red cluster indicates that antennae preferred greater angles (away from odor source). The red cluster gradually disappears as the odor presentation is maintained. Another red, albeit less intense, cluster appears about 1 s after odor conclusion. We name the second red clusters “Rebound Clusters”. Consistent with the mean angle recovery times above, we note that the rebound clusters seem to appear earlier and are less intense for diluted odors than for pure odors.

## DISCUSSION

The method described above enables real-time tracking of insect antenna and proboscis movements without requiring special animal preparations or hardware.

### Limitations

Despite these advantages, there are some limitations of the method. These include the requirement that the head of the animal is restricted from movement, the need for the user to select the location and orientation of the animal for each video, the requirement to have access to a Windows computer, and the software’s inability to track movement in three dimensions and in some visually ambiguous appendage positions described below.

The software requires that the head of the animal is fixed in place and is not moving during the video. This is similar to the preparations of previous work^48–51^. It is possible to modify the software to allow automatic detection of head rotations, however, this would consume additional processing time and introduce a new source of error. If the modified software were to detect the head rotation incorrectly, this would affect the antennae angles, as their computation is relative to the head rotation angle. Currently, the user selects the head orientation once per video. This approach, while not without human error, minimizes angle calculation errors when the head is not allowed to move during the video.

The software also requires a Windows 7 (or later) operating system (OS). Our goal was to make the software easy to install, setup, and use by users without programming or sophisticated computer administration skills. We decided to target Windows because it is widely available, and in cases where access to it is limited, virtual machines (e.g. VirtualBox, VMware, Parallels) with Windows can be easily created. This choice of OS greatly simplifies software installation through the use of an easy-to-use, command-line-free installer and avoids bugs specific to different OSs.

The software only tracks the position of the appendages in 2D space. Insects are known to move their antenna in three dimensions, which could mean that important information is lost when only 2D coordinates are measured. While the use of multiple cameras or mirrors could aid in collecting the additional information required for 3D localization, it is possible to compute, with the use of trigonometric relations, an estimated out-of-plane position by assuming that the antennae are single line segments of constant length and only move on one side of the camera plane. For honey bees, this assumption holds true to obtain rough estimates for the position in the third dimension, but would not necessarily be the case for other species and situations.

The software will not correctly detect the antennae and proboscis tip locations in some ambiguous situations. If an animal moves an antenna so that in the video it overlaps an extended proboscis, the software will likely detect the tip of the antenna as the tip of the proboscis. The antenna angle however, will still likely be computed correctly (from the nonoverlapping part). Similarly, if the antenna tips move directly above the head of the animal (i.e. not on the sides) then the software might only detect the part of the antenna that is visible outside of the head, or assume the previous location of the antenna, or detect spurious video noise as antenna location. In both of the situations, even human raters have difficulty discerning the antenna from the proboscis or the head. To mitigate this problem, we recommend applying a 3-frame, symmetric rolling median^57^ filter to the raw X and Y coordinates produced by the software. This filter removes large transient (single-frame) position fluctuations, and preserves longer antenna position movements. In our tests, 3-frame filter performed better than no filter, while wider filters (e.g. 5, 11, or 15 frames) reduced accuracy. Example R code that uses the filter and a video tutorial can be found online^58^.

### Value as a Scientific Tool

The availability of a method to rapidly obtain accurate insect appendage movements in a cost-effective manner has the potential to open up new areas of investigation.

Proboscis extension reflex (PER) is a commonly used behavioral response to investigate learning and memory of a variety of insects^59^. Previous studies have generally relied on a binary extended-or-not measure of PER, although video and electromyographic analyses have shown much more complex topologies to proboscis movements^65,66^. Our method allows rapid quantification of proboscis movements in high temporal and spatial resolutions.

Insect antenna movements in response to odors are poorly understood. One reason for this is that the antennae tend to move so rapidly that a cost-effective, automated means to obtain antenna movement data has not been available. Our method could be used to rapidly obtain antenna movement data for large numbers of insects in a large number of conditions. This could aid, for example, researchers investigating the mapping between antenna movements in response to various stimuli, in particular volatile odors. Using cameras that capture frames at 30 Hz, the software can be used to characterize antennal movement dynamics up to 15 Hz (Nyquist limit). If characterization in higher frequencies is needed, cameras with higher capture rates (e.g. 60 or 120 fps) could be utilized. However, a faster computer may be required to process higher fps videos in real-time. We speculate that classes of odors, and possibly even some individual odors, have characteristic innate antennal movements. If those classes or compounds could be discovered, unknown odors or their class could be detected from antennal movement of untrained insects. If such a mapping exists, then the combination of sufficient antenna movement data and state of the art machine learning algorithms should begin to uncover it. Also, how that mapping changes in response to learning, forms during development, or is disrupted with genetic interventions could offer insight into functions of the olfactory system. Finally, this work could give insight into artificial detection of odors if it reveals optimal sampling methods for odors in complex environments.

### Future Work

Here, we showed that antenna movement data can be rapidly obtained and analyzed. We showed significant behavior responses can be detected from the data generated by our software, and have identified several areas of further investigation.

The time courses of stimulus-elicited antenna angle deviations from and recovery to baseline and any stimulus-conclusion rebound effects and its dependence on odor concentration can be investigated and modeled mathematically. Additionally, any changes of antenna movements induced by appetitive or aversive conditioning also can be assessed with the software.

Better differentiation of odors can also be explored. In this study, both odors, in pure and 35x diluted versions elicited similar responses: the antennae, on average, appeared to rapidly withdraw away from the odor source and return to pre-odor baselines after a few seconds. We speculate that even the diluted versions may have been very strong olfactory stimuli for the honey bees. If true, a broader range of concentrations could be used to determine if the antennal responses differentiate the odors. Additionally, more sophisticated analysis may better reveal differences in antennal movements in response to different odors. We have made the data files used to generate figures in this manuscript available to interested researchers on the SwarmSight website^67^.

Furthermore, while outside the scope of this manuscript, the software could be extended to process videos of animals placed in chambers with dual mirrors angled at 45° (see Figure 1D for example). This could be used to accurately localize and track the appendages and their movement in 3D space. However, the algorithms for 3D tracking would be required to efficiently a) disambiguate between multiple antennae when they are visible in one of the side mirrors, b) correct for imperfections in mirror angles, and c) account for distortions due to camera positioning.

Finally, additional gains in position accuracy might be realized via the use of a Kalman filter^68^, which models and utilizes physical state information such as appendage velocity and acceleration to constrain predicted locations. However, any gains in accuracy should be evaluated against any reductions in speed due to additional computations.

## Conclusion

Many insects use antennae to actively sample volatile compounds in their local environments. Patterns in antennal movements may provide insight into insect odor perception and how it is affected by conditioning, toxic compounds, and genetic alterations. Similarly, proboscis movements have been used to assess odor perception and its modulation. However, rapidly obtaining large quantities of high-resolution appendage movement data has been difficult. Here, we have described a protocol and software that automates such task. In summary, we have created and demonstrated how the combination of inexpensive hardware, a common animal preparation, and our open-source software can be used to rapidly obtain high-resolution insect appendage movement data. We showed the output of the software, how it outperforms human raters in speed and accuracy, and how its output data can be analyzed and visualized.

## ACKNOWLEDGMENTS

JB, SMC, and RCG were supported by NIH R01MH1006674 to SMC and NIH R01EB021711 to RCG. CMJ and BHS were supported by NSF Ideas lab project on “Cracking the olfactory code” to BHS.

## DISCLOSURES

The authors declare that they have no competing financial interests.

## REFERENCES

1. Hansson B. S. Insect Olfaction. (Springer Science & Business Media: 1999).

2. Masson, C., Mustaparta, H. & others Chemical information processing in the olfactory system of insects. Physiological Reviews 70 (1), 199–245 (1990).

3. Firestein S. How the olfactory system makes sense of scents. Nature 413 (6852), 211–218 (2001).

4. Schneider D. Insect antennae. Annual review of entomology 9 (1), 103–122 (1964).

5. Kaissling K. Chemo-electrical transduction in insect olfactory receptors. Annual review of neuroscience 9 (1), 121–145 (1986).

6. Nakagawa T. & Vosshall L. B. Controversy and consensus: noncanonical signaling mechanisms in the insect olfactory system. Current Opinion in Neurobiology 19 (3), 284–292, doi:10.1016/j.conb.2009.07.015 (2009).

7. Heisenberg M. What do the mushroom bodies do for the insect brain? An introduction. Learning & Memory 5 (1), 1–10 (1998).

8. Zars T. Behavioral functions of the insect mushroom bodies. Current opinion in neurobiology 10 (6), 790–795 (2000).

9. Heisenberg M. Mushroom body memoir: from maps to models. Nature Reviews Neuroscience 4 (4), 266–275 (2003).

10. Pelletier Y. & McLEOD, C. D. Obstacle perception by insect antennae during terrestrial locomotion. Physiological Entomology 19 (4), 360–362 (1994).

11. Suzuki H. Antennal movements induced by odour and central projection of the antennal neurones in the honey-bee. Journal of Insect Physiology 21 (4), 831–847 (1975).

12. Wachowiak M. All in a sniff: olfaction as a model for active sensing. Neuron 71 (6), 962–973 (2011).

13. Bruce, T. J., Wadhams, L. J. & Woodcock, C. M. Insect host location: a volatile situation. Trends in plant science 10 (6), 269–274 (2005).

14. Lunau K. & Wacht S. Optical releasers of the innate proboscis extension in the hoverfly Eristalis tenax L.(Syrphidae, Diptera). Journal of Comparative Physiology A: Neuroethology, Sensory, Neural, and Behavioral Physiology 174 (5), 575–579 (1994).

15. Szucsich N. U. & Krenn H. W. Morphology and function of the proboscis in Bombyliidae (Diptera, Brachycera) and implications for proboscis evolution in Brachycera. Zoomorphology 120 (2), 79–90 (2000).

16. Harder L. D. Measurement and estimation of functional proboscis length in bumblebees (Hymenoptera: Apidae). Canadian Journal of Zoology 60 (5), 1073–1079 (1982).

17. Hobbs G. A. Further studies on the food-gathering behaviour of bumble bees (Hymenoptera: Apidae). The Canadian Entomologist 94 (05), 538–541 (1962).

18. Krenn H. W. Functional morphology and movements of the proboscis of Lepidoptera (Insecta). Zoomorphology 110 (2), 105–114 (1990).

19. Krenn H. W. Feeding mechanisms of adult Lepidoptera: structure, function, and evolution of the mouthparts. Annual review of entomology 55, 307–327 (2010).

20. Hepburn H. R. Proboscis extension and recoil in Lepidoptera. Journal of Insect Physiology 17 (4), 637–656 (1971).

21. Ramírez, G., Fagundez, C., Grosso, J. P., Argibay, P., Arenas, A. & Farina, W. M. Odor Experiences during Preimaginal Stages Cause Behavioral and Neural Plasticity in Adult Honeybees. Frontiers in Behavioral Neuroscience 10, doi:10.3389/fnbeh.2016.00105 (2016).

22. Takeda K. Classical conditioned response in the honey bee. Journal of Insect Physiology 6 (3), 168–179, doi:10.1016/0022-1910(61)90060-9 (1961).

23. Bitterman, M. E., Menzel, R., Fietz, A. & Schäfer, S. Classical conditioning of proboscis extension in honeybees (Apis mellifera). Journal of comparative psychology 97 (2), 107 (1983).

24. Lambin, M., Armengaud, C., Raymond, S. & Gauthier, M. Imidacloprid-induced facilitation of the proboscis extension reflex habituation in the honeybee. Archives of insect biochemistry and physiology 48 (3), 129–134 (2001).

25. Masterman, R., Smith, B. H. & Spivak, M. Brood odor discrimination abilities in hygienic honey bees (Apis mellifera L.) using proboscis extension reflex conditioning. Journal of Insect Behavior 13 (1), 87–101 (2000).

26. Rix R. R. & Christopher Cutler, G. Acute Exposure to Worst-Case Concentrations of Amitraz Does Not Affect Honey Bee Learning, Short-Term Memory, or Hemolymph Octopamine Levels. Journal of Economic Entomology 110 (1), 127–132, doi:10.1093/jee/tow250 (2017).

27. Urlacher E. et al. Measurements of Chlorpyrifos Levels in Forager Bees and Comparison with Levels that Disrupt Honey Bee Odor-Mediated Learning Under Laboratory Conditions. Journal of Chemical Ecology 42 (2), 127–138, doi:10.1007/s10886-016-0672-4 (2016).

28. Charbonneau, L. R., Hillier, N. K., Rogers, R. E. L., Williams, G. R. & Shutler, D. Effects of Nosema apis, N. ceranae, and coinfections on honey bee (Apis mellifera) learning and memory. Scientific Reports 6, doi:10.1038/srep22626 (2016).

29. Urlacher, E., Devaud, J.-M. & Mercer, A. R. C-type allatostatins mimic stress-related effects of alarm pheromone on honey bee learning and memory recall. PLOS ONE 12 (3), e0174321, doi:10.1371/journal.pone.0174321 (2017).

30. Eiri D. M. & Nieh J. C. A nicotinic acetylcholine receptor agonist affects honey bee sucrose responsiveness and decreases waggle dancing. Journal of Experimental Biology 215 (12), 2022–2029, doi:10.1242/jeb.068718 (2012).

31. Liang, C.-H., Chuang, C.-L., Jiang, J.-A. & Yang, E.-C. Magnetic Sensing through the Abdomen of the Honey bee. Scientific Reports 6, doi:10.1038/srep23657 (2016).

32. Erber, J., Pribbenow, B., Bauer, A., Kloppenburg, P. & others Antennal reflexes in the honeybee: tools for studying the nervous system. Apidologie 24, 283–283 (1993).

33. Erber, J., Kierzek, S., Sander, E. & Grandy, K. Tactile learning in the honeybee. Journal of Comparative Physiology A 183 (6), 737–744, doi:10.1007/s003590050296 (1998).

34. Erber J. & Pribbenow B. Antennal Movements in the Honeybee: How Complex Tasks are Solved by a Simple Neuronal System. Prerational Intelligence: Adaptive Behavior and Intelligent Systems Without Symbols and Logic, Volume 1, Volume 2 Prerational Intelligence: Interdisciplinary Perspectives on the Behavior of Natural and Artificial Systems, Volume 3, 109–121, doi:10.1007/978-94-010-0870-9_9 (2000).

35. McAfee, A., Collins, T. F., Madilao, L. L. & Foster, L. J. Odorant cues linked to social immunity induce lateralized antenna stimulation in honey bees (Apis mellifera L.). Scientific Reports 7, doi:10.1038/srep46171 (2017).

36. Dötterl, S., Vater, M., Rupp, T. & Held, A. Ozone Differentially Affects Perception of Plant Volatiles in Western Honey Bees. Journal of Chemical Ecology 42 (6), 486–489, doi:10.1007/s10886-016-0717-8 (2016).

37. Wang Z. et al. Honey Bees Modulate Their Olfactory Learning in the Presence of Hornet Predators and Alarm Component. PLOS ONE 11 (2), e0150399, doi:10.1371/journal.pone.0150399 (2016).

38. Søvik, E., Plath, J. A., Devaud, J.-M. & Barron, A. B. Neuropharmacological Manipulation of Restrained and Free-flying Honey Bees, Apis mellifera. JoVE (Journal of Visualized Experiments) (117), e54695–e54695, doi:10.3791/54695 (2016).

39. Fang, Y., Du, S., Abdoola, R., Djouani, K. & Richards, C. Motion Based Animal Detection in Aerial Videos. Procedia Computer Science 92, 13–17 (2016).

40. Miller, B., Lim, A. N., Heidbreder, A. F. & Black, K. J. An Automated Motion Detection and Reward System for Animal Training. Cureus 7 (12), doi:10.7759/cureus.397

41. Birgiolas, J., Jernigan, C. M., Smith, B. H. & Crook, S. M. SwarmSight: Measuring the temporal progression of animal group activity levels from natural-scene and laboratory videos. Behavior research methods, 1–12 (2016).

42. Stern, U., Zhu, E. Y., He, R. & Yang, C.-H. Long-duration animal tracking in difficult lighting conditions. Scientific reports 5, 10432 (2015).

43. Macrì, S., Mainetti, L., Patrono, L., Pieretti, S., Secco, A. & Sergi, I. A tracking system for laboratory mice to support medical researchers in behavioral analysis. Engineering in Medicine and Biology Society (EMBC), 2015 37th Annual International Conference of the IEEE, 4946–4949at <http://ieeexplore.ieee.org/abstract/document/7319501/> (2015).

44. Crall, J. D., Gravish, N., Mountcastle, A. M. & Combes, S. A. BEEtag: a low-cost, image-based tracking system for the study of animal behavior and locomotion. PloS one 10 (9), e0136487 (2015).

45. York, J. M., Blevins, N. A., McNeil, L. K. & Freund, G. G. Mouse short-and long-term locomotor activity analyzed by video tracking software. JoVE (Journal of Visualized Experiments) (76), e50252–e50252 (2013).

46. Noldus, L. P., Spink, A. J. & Tegelenbosch, R. A. EthoVision: a versatile video tracking system for automation of behavioral experiments. Behavior Research Methods 33 (3), 398–414 (2001).

47. Egnor S. E. R. & Branson K. Computational Analysis of Behavior. Annual Review of Neuroscience 39 (1), 217–236, doi:10.1146/annurev-neuro-070815-013845 (2016).

48. Cholé, H., Junca, P. & Sandoz, J.-C. Appetitive but not aversive olfactory conditioning modifies antennal movements in honeybees. Learning & Memory 22 (12), 604–616 (2015).

49. Shen M. et al. Interactive tracking of insect posture. Pattern Recognition 48 (11), 3560–3571, doi:10.1016/j.patcog.2015.05.011 (2015).

50. Mujagić, S., Würth, S. M., Hellbach, S. & Dürr, V. Tactile conditioning and movement analysis of antennal sampling strategies in honey bees (apis mellifera l.). Journal of visualized experiments: JoVE (70)at <https://www.ncbi.nlm.nih.gov/pmc/articles/PMC3578280/> (2012).

51. Shen, M., Szyszka, P., Deussen, O., Galizia, C. G. & Merhof, D. Automated tracking and analysis of behavior in restrained insects. Journal of neuroscience methods 239, 194–205 (2015).

52. Bellard, F., Niedermayer, M. & others FFmpeg. Availabel from: http://ffmpeg.org (2012).

53. Cucchiara, R., Grana, C., Piccardi, M. & Prati, A. Detecting moving objects, ghosts, and shadows in video streams. IEEE transactions on pattern analysis and machine intelligence 25 (10), 1337–1342 (2003).

54. Shaw J. R. QuickFill: An efficient flood fill algorithm. The Code Project (2004).

55. Zhang H. & Wu, K. A vehicle detection algorithm based on three-frame differencing and background subtraction. Computational Intelligence and Design (ISCID), 2012 Fifth International Symposium on 1, 148–151 at <http://ieeexplore.ieee.org/abstract/document/6406940/> (2012).

56. Elgammal, A., Harwood, D. & Davis, L. Non-parametric Model for Background Subtraction. Computer Vision — ECCV 2000, 751–767, doi:10.1007/3-540-45053-X_48 (2000).

57. Zeileis A. & Grothendieck G. zoo: S3 infrastructure for regular and irregular time series. arXiv preprint math/0505527 at <https://arxiv.org/abs/math/0505527> (2005).

58. SwarmSight.org. at <http://swarmsight.org/>

59. Smith B. H. & Burden C. M. A Proboscis Extension Response Protocol for Investigating Behavioral Plasticity in Insects: Application to Basic, Biomedical, and Agricultural Research. JoVE (Journal of Visualized Experiments) (91), e51057–e51057, doi:10.3791/51057 (2014).

60. How to split a video or audio file with VLC Player. Darktips at <https://darktips.com/split-video-audio-file/> (2012).

61. SwarmSight Appendage Tracking CSV File Column Reference | SwarmSight by JustasB. at <http://swarmsight.org/Examples/Appendage%20Tracking/ColumnReference>

62. R Core Team R: A Language and Environment for Statistical Computing. at <http://www.R-project.org/> (R Foundation for Statistical Computing: Vienna, Austria, 2014).

63. Team, Rs. RStudio: integrated development for R. RStudio, Inc., Boston, MA URL http://www.rstudio.com (2015).

64. Venables W. N. & Ripley B. D. Modern Applied Statistics with S-PLUS. (Springer Science & Business Media: 2013).

65. Smith B. H. & Menzel R. An Analysis of Variability in the Feeding Motor Program of the Honey Bee; the Role of Learning in Releasing a Modal Action Pattern. Ethology 82 (1), 68–81, doi:10.1111/j.1439-0310.1989.tb00488.x (1989).

66. Smith B. H. An analysis of blocking in odorant mixtures: an increase but not a decrease in intensity of reinforcement produces unblocking. Behavioral neuroscience 111 (1), 57 (1997).

67. Birgiolas J. SwarmSight Antenna Tracking CSV files. at <https://github.com/JustasB/SwarmSight/tree/master/Examples/Appendage%20Tracking/Birgiolas%20et.%20al.%20(2015)%20JOVE%20figures/Figures%204%265/CSVs>

68. Kalman R. E. A New Approach to Linear Filtering and Prediction Problems. Journal of Basic Engineering 82 (1), 35–45, doi:10.1115/1.3662552 (1960).

